# Characterization of Ovine Abortion in Uruguay Reveals Extensive Non-Clonal Diversity and Multiple Evolutionary Origins of *Toxoplasma gondii*

**DOI:** 10.64898/2026.03.31.715541

**Authors:** Leandro Tana-Hernandez, Pablo Fresia, Andrés M. Cabrera, Alejandra Valentin, Matías Dorsch, Sergio Fierro, Federico Giannitti, Luisa Berná, María E. Francia

## Abstract

*Toxoplasma gondii* is a globally prevalent zoonotic parasite with multiple life stages and transmission routes, including ingestion and transplacental transmission. It is a major cause of abortion in sheep, goats and pigs, among other production animals, worldwide. While Type II strains are common in livestock in North America and Europe, non-archetypal, non-clonal genotypes are highly prevalent in South America. This study aimed to determine the molecular epidemiology of *T. gondii* strains causing sheep abortion in Uruguay. Phylogenomic analyses confirmed significant divergence among typed strains and revealed similarities with genotypes previously detected in the human population. Two novel strains, were isolated and characterized, uncovering the connection between their genetic makeup and phenotypes. Differences in virulence could be correlated to differences in gene copy number of the pseudo kinase ROP5 – further highlighting this virulence factor as relevant in wild strains. Whole-genome sequencing further confirmed the divergence among Uruguayan isolates, uncovering at least three distinct evolutionary origins. Overall, our findings highlight the circulation of virulent non-clonal lineages with links to human infections and underscore the importance of furthering genomic surveillance in South America to better understand *Toxoplasma*’s transmission dynamics, pathogenic potential, and zoonotic risk.

## INTRODUCTION

*Toxoplasma gondii* is a zoonotic protozoan parasite of worldwide distribution, which showcases its ability to infect most warm-blooded vertebrates, including terrestrial and marine mammals, and birds [1,2]. Its success as a parasite is partially owed as well to its multiple life stages, and multiple transmission routes. Within the intermediate hosts, *T. gondii* can exist as a fast replicating tachyzoite capable of invading and destroying virtually any nucleated cell, causing an acute infection. It can then persist chronically in a latent encysted form lodging bradyzoite[3]. Cats and other felids serve as definitive hosts sustaining parasite sexual stages, which enables sexual recombination, and the release of infectious oocysts into the environment [4,5]. Transmission occurs through ingestion of food contaminated with sporulated oocysts or raw, undercooked meat that contains bradyzoite tissue cysts [6]. Notably, *T. gondii* can also be transmitted transplacentally, which leads to congenital infection and/or abortion [7]. Though the impact of transplacental transmission of *T. gondii* on human abortion has not been systematically quantified, it is well established that certain economically relevant species such as sheep, goat and pigs, are particularly sensitive to *T. gondii* infection during pregnancy. *T. gondii* is in fact a leading cause of reproductive loss in these species [8].

The population genetic structure of *T. gondii* both in North America and Europe is primarily composed of clonal lineages originally termed “typical.” These genetic types, named I through III, differ in virulence and species distribution, with type I strains being the rarest in nature yet the most virulent, and type II strains occurring most frequently and displaying intermediate virulence [9,10]. Type II strains are also the most frequently found in sheep and other livestock [11,12]. On the other hand, it is well established that *T. gondii* non-clonal genotypes are widely prevalent in South America [13]. These genotypes arise most frequently as combinations of clonal alleles found in strain of type I-III strains but also include novel non-clonal alleles; these strains are collectively known as non-archetypal. Even though the relationship between non-clonal genotypes and phenotype is yet to be fully clarified, it is widely accepted that non clonal genotypes are frequently associated with phenotypic traits of exacerbated virulence, tendency to encyst and drug resistance [14–16]. Most research aiming to uncover the molecular underpinnings of phenotypes has relied on laboratory-adapted strains. These studies have identified several genes related to virulence. In particular, gene copy number variations of the secreted pseudo kinase ROP5 has been shown to play a pivotal role; the higher the number of ROP5 genes, the greater the virulence of the strain [17,18]. However, many of these extensively studied strains have lost key virulence-related traits, such as the ability to form brain cysts in mice, and none display natural drug resistance. Consequently, there is a significant knowledge gap in the understanding of the impact of naturally occurring variability on clinically relevant phenotypes.

Pioneering work from Freyre and colleagues isolated a *T. gondii strain from a* sheep aborted fetus in Uruguay [19,20]. This particular strain, known as “CASTELLS”, belongs to clade E together with other non-archetypal isolates, and exhibits high virulence in primary infections [15,21]. Previous work by our group and others have shown that *T. gondii* is a major cause of abortion in sheep from Uruguay [19,22–24], and other South American countries [25].

Given the high incidence of *T. gondii* in sheep abortions, and the prior isolation of a highly virulent non-clonal strain in the country [20] we set to determine the contribution of distinct *T. gondii* genotypes to the molecular epidemiology of sheep abortion in Uruguay.

Furthermore, to delve into the correlation between non clonal genotypes and their phenotypic implications, we isolate and characterize two novel *T. gondii* strains and finely analyze their genomic structure relating their genetic variability to regional strains and preliminarily identifying the molecular underpinning of their distinct phenotypic traits.

## MATERIALS AND METHODS

### Study samples

Placental and fetal tissue samples were derived from 58 sheep abortions obtained from sheep flocks residing in different locations within Uruguay (as indicated in **Supplementary Figure 1**). Samples were voluntarily rendered by sheep owners and first sent to the “Plataforma de Investigación en Salud Animal, Estación Experimental INIA La Estanzuela” in Colonia, Uruguay for diagnosis. Here, *T. gondii* infection and associated disease was confirmed by histopathology, and positive parasite DNA detection by PCR as reported previously [22]. All samples were reanalyzed for *T. gondii* DNA detection as described below prior to proceeding with the *in silico* RFLP analysis.

### DNA extraction and Identification of *Toxoplasma gondii*

DNA from tissue or cell culture, was extracted using the Quick-DNA Tissue/Insect MiniPrep Kit (D6016, Zymo Research) following the supplier’s recommendations. DNA was stored at - 20°C or 4°C until further analysis *T. gondii* DNA detection was carried out by polymerase chain reaction (PCR) using specific primers specified in Table S1, and a MangoMix master mix (BIO-25033, Meridian Biosciences) following the supplier’s recommendations. PCR detection of *T. gondii* DNA was performed as described in [26] and [27] without modification.

### *In silico* PCR-RFLP genotyping of *Toxoplasma gondii*

Strain genotype determination (also referred to as “typing”) was carried out by in *silico* analyses of genetic markers as previously described by [28]. Amplification of nine markers (SAG2, SAG3, BTUB, GRA6, c22-8, c-29-2, L358, PK1 and Apico) was pursued by PCR using the oligonucleotides and cycling parameters listed in Table S1. We note that out of the nine polymorphic markers selected for typing, we were consistently able to amplify all but the “Apico”. Following successful amplification of markers, Sanger sequencing of amplicons was performed (Macrogen Inc., South Korea). Detection of polymorphic profiles was carried out as described by Castro and collaborators [29] and used as in [30] without modifications. Genotype assignment is outlined by strain type in **Supplementary Table 1**. All marker sequences obtained from the analyzed samples were submitted to GenBank, and the corresponding accession numbers are detailed in **Supplementary Table 2**.

### Phylogenetic analysis of ovine abortion samples

To evaluate the phylogenetic relations among *T. gondii* strains present in abortion-derived samples, types by *in silico* RFLP analyses, we used the same strategic methodology as in [30]. Briefly, we first visually inspected the sequencing chromatograms and manually curated overlapping peaks and erroneously called bases. The curated sequences for each molecular marker of the amplified isolates were aligned using the MAFFT package [31,32] with the ‘L-INS-i’ method and parameters ‘--localpair’ and ‘--maxiterate = 1,000’. Alignment confidence scores were computed with rGUIDANCE [33] and the resulting confidence weights were incorporated into the analysis via a weights file.

A maximum likelihood (ML) tree was constructed in RAxML using the GTRCAT substitution model. A Neighbor-Joining (NJ) tree based on the TN93 distance served as the starting topology. Gene-partitioned analyses were specified through a partition file (-q partitions), and the final tree was generated using an exhaustive search (-f I). To evaluate node support, 100 bootstrap replicates were performed with the rapid bootstrap algorithm (-f a), and the resulting support values were mapped onto the ML tree.

### Use of animals for experimentation

All protocols pertaining to the use of experimental animals were pre-approved by the Institutional Ethics Committee of the Institut Pasteur de Montevideo (CEUA No. 019-19 and 023-22). Animals were bred at the Laboratory Animal Biotechnology Unit of the Institut Pasteur de Montevideo under specific pathogen-free conditions in ventilated racks (IVC, 1285L, Tecniplast, Milan, Italy), housed in pathogen-free specific facilities and subjected to daily monitoring.

Humanitarian criteria for animal endpoints were strictly followed in accordance with the established norms. Euthanasia was performed by first inducing deep anesthesia through intraperitoneal administration of a ketamine/xylazine combination. Mice were anesthetized by intraperitoneal injection of a freshly prepared ketamine–xylazine 2% solution, administered at a volume of 10 mL/kg, corresponding to doses of 110 mg/kg ketamine (phs® Pharmaservice Montevideo-Uruguay) and 13 mg/kg xylazine (Laboratorios Microsules Uruguay S.A.). Anesthetic depth was verified by the absence of motor responses to nociceptive stimulation of the forelimbs and hindlimbs, after which cervical dislocation was carried out as the physical method of euthanasia. All experiments were carried out in accordance with relevant guidelines outlined in the animal experimentation protocol, abiding by the national animal protection law (No. 18.611) and the ARRIVE guidelines[34].

### Strain Isolation

*S*trains isolation was carried out as described in [35] with modifications. In short, placenta from a fetal loss case derived in the Department of Colonia in 2021 (TgUru2) or fetal brain (TgUru1) from an aborted fetus from Florida were used. Tissues were homogenized in Phosphate-buffered saline (PBS) solution containing antibiotics using a manual douncer. DNA was extracted from a fraction of the homogenate to determine by PCR whether the sample contained traces of *T. gondii* DNA, following the PCR-based DNA detection procedures described above. A maximum of 200 µl of PCR-positive homogenates were intraperitoneally injected into Interferon gamma knockout B6 mice. Mice were evaluated daily to detect signs of toxoplasmosis. Mice exhibiting symptoms such as weight loss, or neurological signs were euthanized. Brain samples and peritoneal fluid were obtained, analyzed by PCR and phase contrast microscopy for the identification of *T. gondii*, and subsequently inoculated and maintained in VERO cells.

### Parasite and cell culture

Parasites were maintained by serial passage in VERO cells cultured in Dubelco’s modified eagle medium (DMEM; Capricorn Scientific GmBH) supplemented with a 1% penicillin/streptomycin (GIBCO, USA) solution and 10% fetal bovine serum (GIBCO, Brazil), at 37C and a 5% CO_2_.

### *In vivo* phenotypic characterization

To assess the virulence of the isolates during the acute phase, 4 6 to 8-week old female BALB/c mice per group, with an average weight of 17,6g at day 0 of infection, were intraperitoneally infected with the indicated parasite doses of each isolate in PBS. In all cases, sterile PBS was included as negative control. Seroconversion was systematically evaluated in surviving mice by Western blot (data not shown). Mice were monitored daily throughout the course of the infection.

### Invasion and growth assays

Intracellular growth and invasion assays were conducted following the procedures described by [36] with the following modifications; assays were performed by infecting confluent neonatal Human Dermal Fibroblasts, (HDFn) cultured on 13 mm coverslips with a total of 1000 parasites. Quantifications were performed by determining the number of parasites per field (invasion) and the number of parasites per vacuole 24 hours post-infection (growth). Parasites were visualized by immunofluorescence assays performed as previously described by [37], using an Olympus epifluorescence microscope and a 60x oil lens. All assays were conducted in triplicate, with the *Toxoplasma gondii* PruΔKu80 strain serving as control, and 50 randomly selected fields were quantified for each experiment.

### Brain cyst counts

BALB/c mice were infected with 100 parasites each with either TgPruΔKu80 or TgUru2. Brain cysts were counted 15 days post-infection following the protocol described by [38]. Briefly, brain tissue was extracted from euthanized mice and divided into two parts, one part was stored at -80°C for further analysis while the other was homogenized using a Pellet Pestle, fixed with 3% formaldehyde for 20 minutes, quenched with 0.1 mM glycine solution for 3 minutes, and blocked/permeabilized overnight at 4°C in blocking solution (3% BSA, 0.2% Triton X-100).

Permeabilized homogenates were stained with DBA conjugated to Fluorescein (Vector Laboratories) (1:500) for 1 h at room temperature. 5 μl of each sample were placed on glass slides and coverslips in triplicate. Cyst counts were pursued using an Olympus IX81 epifluorescence microscope using a 40X objective. Plotting and Statistical analysis were carried out using the GraphPad. Comparisons were analysed using a student’s unpaired t-test.

### Whole-genome sequencing

DNA was extracted as described above. Whole-genome sequencing was carried out using two complementary approaches: long-read sequencing with Oxford Nanopore Technologies (ONT) and short-read sequencing with the DNBseq platform (BGI Genomics, Hong Kong). For ONT, libraries were prepared with the Rapid Barcoding Kit (SQK-RBK004, ONT) from 400 ng of total genomic DNA, following the manufacturer’s instructions, and sequenced for 24 h on a MinION Mk1C device with a Spot-ON Flow Cell R9.4.1 (FLO-MIN106D, ONT). Base calling was performed with Guppy v3.0.3 (ONT). Short-read sequencing was performed in paired-end mode (2 × 150 bp) on the DNBseq platform.

### Genome Assembly, Polishing, and Annotation

For both *T. gondii* strains (TgUru1 and TgUru2), Oxford Nanopore long reads were assembled de novo using Canu [39]. Contigs corresponding to the apicoplast and mitochondrial genomes were identified based on GC content and sequence similarity and were removed from the final assemblies. Contigs with no detectable homology to the *T. gondii* reference genome were manually inspected using BLASTn and found to correspond to non-target contaminant sequences (e.g., *Macaca mulatta* derived from the host cell); these were subsequently discarded.

Assembly polishing was carried out in two successive steps. First, long reads were aligned with minimap2 and used as input for Racon to correct indels and structural errors. Then, paired-end DNBseq short reads (BGI) were aligned using BWA-MEM [40] and two rounds of Pilon [41] were performed to correct base-level mismatches and refine the assemblies.

The resulting polished assemblies were scaffolded against the *T. gondii* RHΔKu80 reference genome -previously assembled by our group- using ABACAS [42] to generate chromosome-level pseudomolecules.

For gene annotation, available RNA-Seq reads (SRR23391969, SRR23391970, SRR23391971) were mapped to the genome assemblies using HISAT2[43,44], and transcript structures were inferred with StringTie [45]. These transcript models were then used to guide gene prediction with the COMPANION pipeline [46], employing *T. gondii* ME49 as the reference genome and default parameters.

For comparative genome analysis, assemblies were aligned using nucmer (part of the MUMmer package [47], and structural variants were identified with Assemblytics [48]. Genomic synteny and rearrangements were visualized using Circos [49].

### Assessment of Copy Number Variation

We examined copy number variation across a curated set of 53 genes encoding rhoptry and dense-granule effectors involved in host invasion and immune modulation. These genes were selected from ToxoDB (gene IDs listed in Supplementary Table 4). Homology searches were performed using BLASTn and BLASTp against the TgUru1 and TgUru2 assemblies to determine gene presence, absence, and copy number relative to the RH reference strain.

Given the differences observed in the number of ROP5 copies among the analysed isolates, we conducted additional analyses to characterize the ROP5 subtypes present in each genome. To this end, the predicted peptide sequences from TgUru1 and TgUru2 were aligned together with all ROP5 protein sequences available in UniProt, totaling 52 sequences (listed in **Supplementary Table 7**). Protein sequences were aligned using MAFFT[32] with the --auto option. Phylogenetic classification of ROP5 variants was carried out using IQ-TREE2[50] under the GTR+F substitution model with 1,000 ultrafast bootstrap and 1,000 SH-aLRT replicates.

To visualize conserved features of the pseudokinase domain, we generated an additional alignment of a representative subset of ROP5 proteins using MAFFT (--auto). This subset included the following UniProt/TrEMBL accessions: I6YX69 (ROP5A), F2YGR7 (ROP5B), I6ZQR7 (ROP5C), F2YGS5, F2YGS7, F5XVD7, Q3YJR4, A0A125YQ30, F5XVD5, and F5XVD6, along with the eight ROP5 copies of TgUru1 and TgUru2.

#### Phylogenetic analysis of strains

To investigate the phylogenetic relationships among isolates, we analyzed a set of 51 strains. This dataset included 47 isolates with predominant ancestry in the American continent with reported hybrid genomes according to [51], together with TgUru1 and TgUru2, and the reference strains TgME49 and TgPru.

Illumina paired end reads for all publicly available isolates were obtained from the NCBI Sequence Read Archive. Accession numbers are listed in **Supplementary Table 6**. Reads were aligned to the *T. gondii* RH reference genome (assembly TgRH_pasteur, GCA_033216535.1 [52]) using BWA-MEM [40]. PCR duplicates were marked with Picard, and BAM files were indexed with SAMtools. Variant calling followed GATK Best Practices: BaseRecalibrator and ApplyBQSR[53] were applied prior to variant discovery with HaplotypeCaller. SNPs and indels were extracted separately using SelectVariants and filtered with hard thresholds. SNPs were retained when they passed the filter QD < 0.2 || QUAL < 30 || SOR > 3.0 || MQRankSum < –12.5 || ReadPosRankSum < –8.0 || FS > 60.0 || MQ < 40.0, while indels were filtered using QD < 0.2 || QUAL < 30 || FS > 200.0 || ReadPosRankSum < – 8.0. Variants failing these criteria were excluded, and high-confidence SNPs and indels were merged per isolate using MergeVcfs [53].

For phylogenetic inference, we focused on a curated set of 100 single-copy nuclear genes distributed across all chromosomes, including several located on unassigned contigs. (listed in **Supplementary Table 5**). For each isolate, gene-specific FASTA sequences were generated by incorporating strain-specific variants using GATK FastaAlternateReferenceMaker [53] These sequences were concatenated into a single multi-locus alignment per isolate, aligned with MAFFT (--auto), and used to infer a maximum-likelihood phylogeny with IQ-TREE2 employing the GTR+F substitution model [50] (ultrafast bootstrap -B 1000 and SH-aLRT -alrt 1000). The resulting tree was visualized using iTOL [54].

## RESULTS

Seventeen sheep samples, labeled TgShUru1-17, exhibiting a positive PCR result for *Toxoplasma gondii* DNA detection were subjected to amplification of 9 conventional genotyping markers. Amplicons were sequenced, and their restriction pattern using established enzymes was analyzed *in silico* **(Figure 1A)**. All 17 samples displayed amplification of at least one typing marker with most samples amplifying Gra6 (14/17) and/or Btub (11/17). PCR amplification success of the remaining markers (SAG2, c29-2, c22-8, L358, PK1, and Apico) was variable.

**FIGURE 1.**
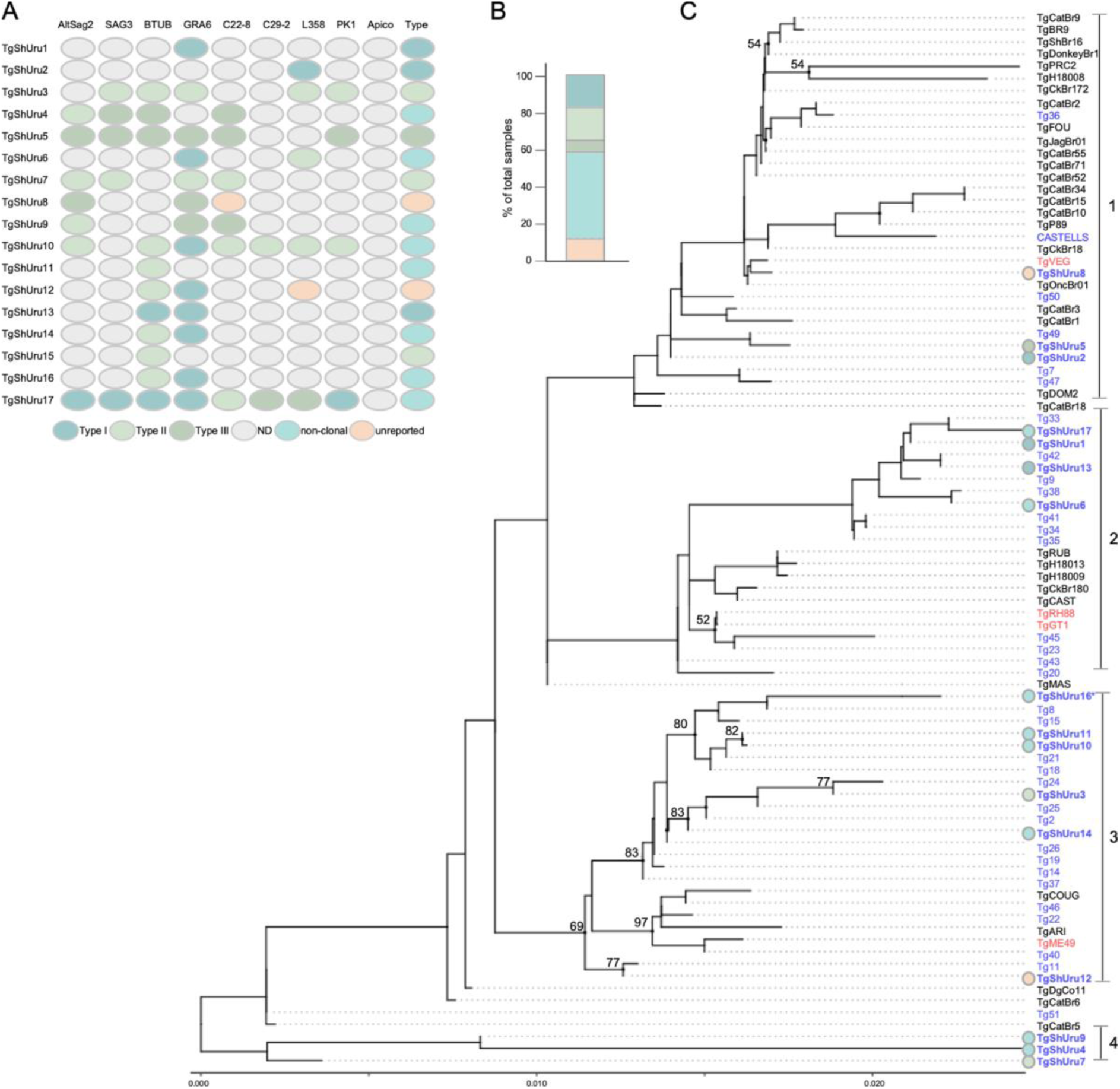
*Genotyping of Toxoplasma gondii* strains causing abortion in sheep. **A. Schematic representation of the in silico RFLP analysis of 17 sheep abortion samples using the indicated markers.** Genetic types were determined by analyzing the combination of genetic types obtained per marker, per sample. Results were color-coded as indicated. ND stands for *not determined*. Note that they “type” column summarizes genotype assignment per strain. **B. Quantification of genetic types found.** The percentage of each assigned genotype in A was plotted. Results are color-coded as in A. **C. Phylogenetic analysis.** A maximum likelihood tree reveals the presence of four phylogenetically distinct groups named 1 through 4. Uruguayan strains are highlighted in blue, and a color-coded dot corresponding to the sample’s genetic type, as identified in A, is shown to the left of each sample for reference. Genetically typed human samples from Uruguay were included and are labeled Tg1-51[30]. Reference strains are highlighted in red. *denotes overlapping position with TgShUru15

Among all characterized samples, 7 samples displayed amplification solely of clonal alleles. Samples TgShUru1, 2 and 13, displayed only type I markers. However, in these samples, only one or two markers could be characterized, indicating that their genotyping was incomplete and highly biased. In contrast, one sample which classified as type III was supported by the amplification of 6 out of the 9 markers (TgShUru5). Finally, three samples classified as type II (TgShUru3, 7, and 15) were supported by the amplification of 5, 4, and 1 out of 9 markers, respectively.

In addition, 8 samples (47%) showed amplification of mixed clonal alleles, indicating the presence of non-clonal hybrid genotypes (**Figure 1B**). In most cases, these corresponded to a mixture of type I and type II (TgShUru6, 10, 11, 14, and 16). However, two samples also showed a mixture of type II and III (TgShUru4 and 9), while one sample displayed a mixture of all three genotypes (TgShUru17). Finally, two samples (TgShUru8 and 12) displayed amplification of non-clonal, previously unreported, alleles for C22-8 and L358, respectively, and were classified as “unreported.”

TgShUru10 and 17 resulted in the greatest number of amplified loci (**Figure 1A**), which allowed us to compare them with reported genotypes in ToxoDB. The genotype detected in sample 10 did not coincide with previously reported genotypes. However, the genotype detected for TgShUru17 matched a previously reported genotype found in samples TgCKGa04 and TgSKMs3. These samples were typed in DNA obtained from a chicken in Guatemala, and a striped skunk in the USA, respectively. No identifying genotype number has been assigned to these samples. Remarkably, out of the 17 analyzed samples, and even with many samples displaying poor amplification, 8 distinct genotypes could be unequivocally distinguished in this study.

To explore how the genetic diversity found differed among strains causing abortion in sheep, we pursued comparative analyses using the complete sequence of amplified alleles (**Figure 1C**). These analyses consider polymorphisms beyond the restriction sites, further resolving existing variability among strains, and allowing comparisons to both reference archetypal strains and publicly available sequences of regional non-archetypal strains. Maximum likelihood analysis resolved four distinct groups of strains (referred to as 1 thru 4). All groups contained genotypes displaying a mix of archetypical alleles. Group 1 contained TgShUru2, 5 and 8, as well as CASTELLS, a strain previously isolated from sheep in Uruguay [23], as well as the TgVEG type III reference strain. Group 2 contained TgShUru1, 6, 13 and 16, as well as the clonal genotypes of TgRH and TgGT1. Group 3 is made up almost exclusively by Uruguayan sequenced genotypes, but also includes TgCOUG, TgARI and the clonal TgME49 Type II strain. Finally, Group 4 includes TgShUru4, 7 and 9, which do not cluster with previously reported sequence types. Interestingly, all amplified loci of TgShUru7 were deemed as type II by *in silico* RFLP, however, this strain clusters away from Type II reference strains when other polymorphisms are considered.

To determine how the *T. gondii* genotypes present in sheep relate to those found in humans, we included in the analysis 32 genotypes previously identified in humans [30]. Conspicuously, sequence types obtained from human patients infected with *T. gondii* from Uruguay are interspersed in groups 1-3 and closely relate to the genotypes found in sheep.

To gauge how non-clonal genotypes translate into phenotypic diversity, we set out to isolate circulating wild strains. For this, we recovered placental tissue from a ewe that had undergone a spontaneous abortion and from the brain of an aborted fetus. Tissue homogenate was inoculated into immuno-naive mice and peritoneal fluid was collected, analyzed by PCR for species confirmation, and transferred to cell culture (**Figures 2A and C, and Supplementary Figure 2**). This led to the isolation of two strains; named TgUru1 and TgUru2. *In silico* RFLP analyses of the 9 typing markers pursued for abortion-derived samples revealed two previously unidentified non-clonal genotypes (**Figure 2A**). In both cases genotyping of the isolated strain coincided with the genotype detected in the originating tissue. TgUru1 predominantly exhibits markers belonging to the type I lineage, in combination with markers that do not fit into any of the clonal classifications and have not been reported previously. TgUru2 predominantly displays markers belonging to type III and type I clonal lineages, and an additional two previously undescribed markers (**Figure 2B**). Comparison to the genotypes available in ToxoDB confirmed that they do not match neither previously genotyped samples nor the previously genotyped strains. Hence, TgUru1 and TgUru2 represent two novel non-clonal strains.

**FIGURE 2.**
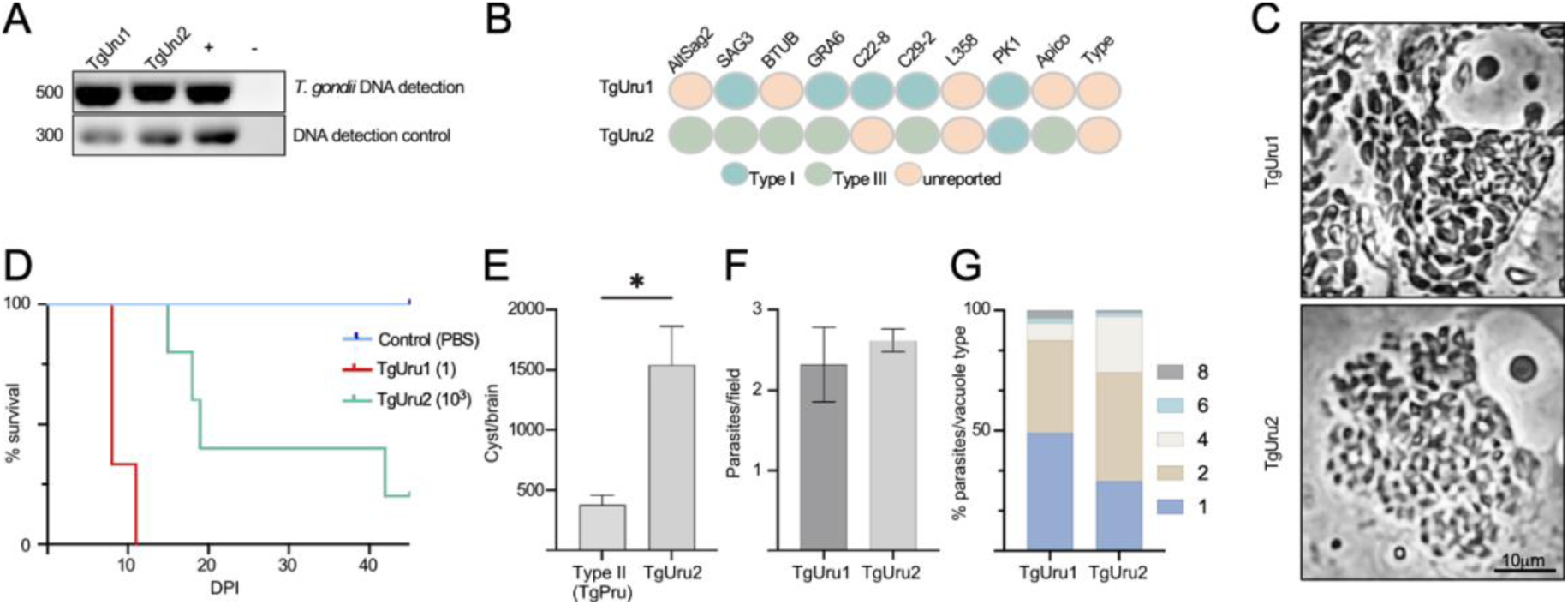
Isolation of two novel non-clonal *Toxoplasma gondii* strains. **A. Isolate identity confirmation by specie-specific PCR**. Detection of *T. gondii* DNA in peritoneal exudates from mice inoculated with homogenates of infected tissue enabled the establishment of two independent isolates named TgUru1 and TgUru2. + denotes positive DNA control. – denotes negative (no DNA) amplification control. Please note that the original gel picture is included in Supplementary Figure 2B. **B**. **Schematic representation of the *in silico* RFLP genotype determination of TgUru1 and TgUru2.** PCR-RFLP genotyping revealed that both isolates are previously unreported non-clonal strains. TgUru1 carries predominantly type I alleles and some unclassified variants, whereas TgUru2 displays a mosaic of type III and type I alleles. No matching genotypes were found in ToxoDB. Results were color-coded as indicated. Note that they “type” column summarizes genotype assignment for each strain. **C**. ***In vitro* phenotype observation**. TgUru1 presents fast and disseminated *in vitro* growth, while TgUru2 forms densely packed vacuoles. Scale bar represents 10μm. **D. *In vivo* virulence determination.** BALB/c (4 animals per condition) were inoculated with the indicated doses of TgUru1 and TgUru2, and with PBS as control. **E. Chronic infection cyst burden quantification**. Surviving TgUru2 infected mice developed a markedly higher cyst load than animals infected with type II strain (TgPRU, mean 1543 vs. 381 cysts/brain; p = 0.0244). **F. Quantification of Invasion per strain**. Both strains invaded host cells at comparable rates, as quantified by vacuole counts at 24 hours post infection. **G. Quantification of Early intracellular growth**. TgUru1 more frequently accumulated in parasite vacuoles of 8, while TgUru2 parasite vacuoles consisted more often of 2-4 parasites, consistent with slower intracellular replication.

We next explored how these strain’s genotypes correlated with physiologically relevant phenotypes. Firstly, we evaluated their *in vivo* virulence in Balb/c mice. Clinical signs consistent with toxoplasmosis (hirsute hair, conjunctivitis, deterioration, and tachypnea) were readily observed in all mice infected with 1 parasite of the TgUru1 strain as early as 7 days post infection, while mice infected with TgUru2 displayed a distinct disease evolution. All mice infected with 10^3^ TgUru2 displayed an asymptomatic infection and 100% survival for the first four weeks post infection. However, 50% of these mice later succumbed to infection by 32 dpi (**Figure 2D**). Therefore, while TgUru1’s lethal dose (LD100) is 1 parasite, TgUru2 displayed a comparable virulence rate to that displayed by the typical Type II strain Pru, with an LD75 of 10^3^ parasites. However, TgUru2 triggered the onset of symptoms comparably later (**Figure 2B**). To further explore TgUru2’s infection kinetics, we asked whether surviving mice were chronically infected. Indeed, we determined that the number of brain cysts generated by TgUru2 infected surviving mice was 4 times higher than those infected with equal doses of TgPru (p=0.0244); while the brains of TgUru2 infected mice averaged 1543 cysts, TgPru infected mice averaged 381 cyst/brain (**Figure 2E**).

To tackle the biology underlying these phenotypes we explored the strain’s *in vitro* behavior. In addition to their detectable genetic differences, TgUru1 and TgUru2 differ dramatically in their *in vitro* growth patterns, with TgUru1 establishing high multiplicity of infection of the host cell cytosol and TgUru2 growing predominantly in clustered rosettes (**Figure 2C**). We quantitated each strain’s capacity to invade and proliferate within host cells. For this, equal numbers of TgUru1 and TgUru2 were allowed to invade, and invaded parasites were allowed to grow for 24 hours. The number of vacuoles per field of view were considered as a *proxy* for the number of invading parasites per strain, while the number of parasites per vacuole were considered a *proxy* for their intracellular replication rate. On average, we found that TgUru1 and TgUru2 exhibited similar rates of invasion as no significant differences between the two were observed (**Figure 2F**). However, we found that TgUru2 exhibited half the number of vacuoles containing 8 parasites compared to TgUru1, with a concomitant increase in the proportion of vacuoles containing 2 and 4 parasites (**Figure 2G**), suggesting that while the strains invade their host cells at comparable rates, once inside the cell, TgUru2’s growth lags.

To investigate the genomic basis of the phenotypes observed in our isolates, we analyzed the structure, variability, and evolutionary relationships of the Uruguayan strains. Both genomes were sequenced using a combination of Oxford Nanopore Technologies long reads and DNBseq short reads, enabling high-contiguity assemblies and accurate polishing. Using publicly available RNA-seq datasets, we generated *de novo* gene annotations for each genome. Summary statistics for the assembled genomes and their corresponding annotations are presented in **Supplementary Table 3**.

Comparative analyses revealed that the overall genomic structure of TgUru1 and TgUru2 closely resembled that of the TgRH reference assembly previously generated by our group, with no major structural rearrangements detected (**Figure 3A** and **Supplementary Figure 3**). Given this structural conservation, we scaffolded both genomes onto the RH reference using ABACAS, recovering 13 chromosomes with lengths and composition comparable to the reference (**Figure 3A**).

**FIGURE 3.**
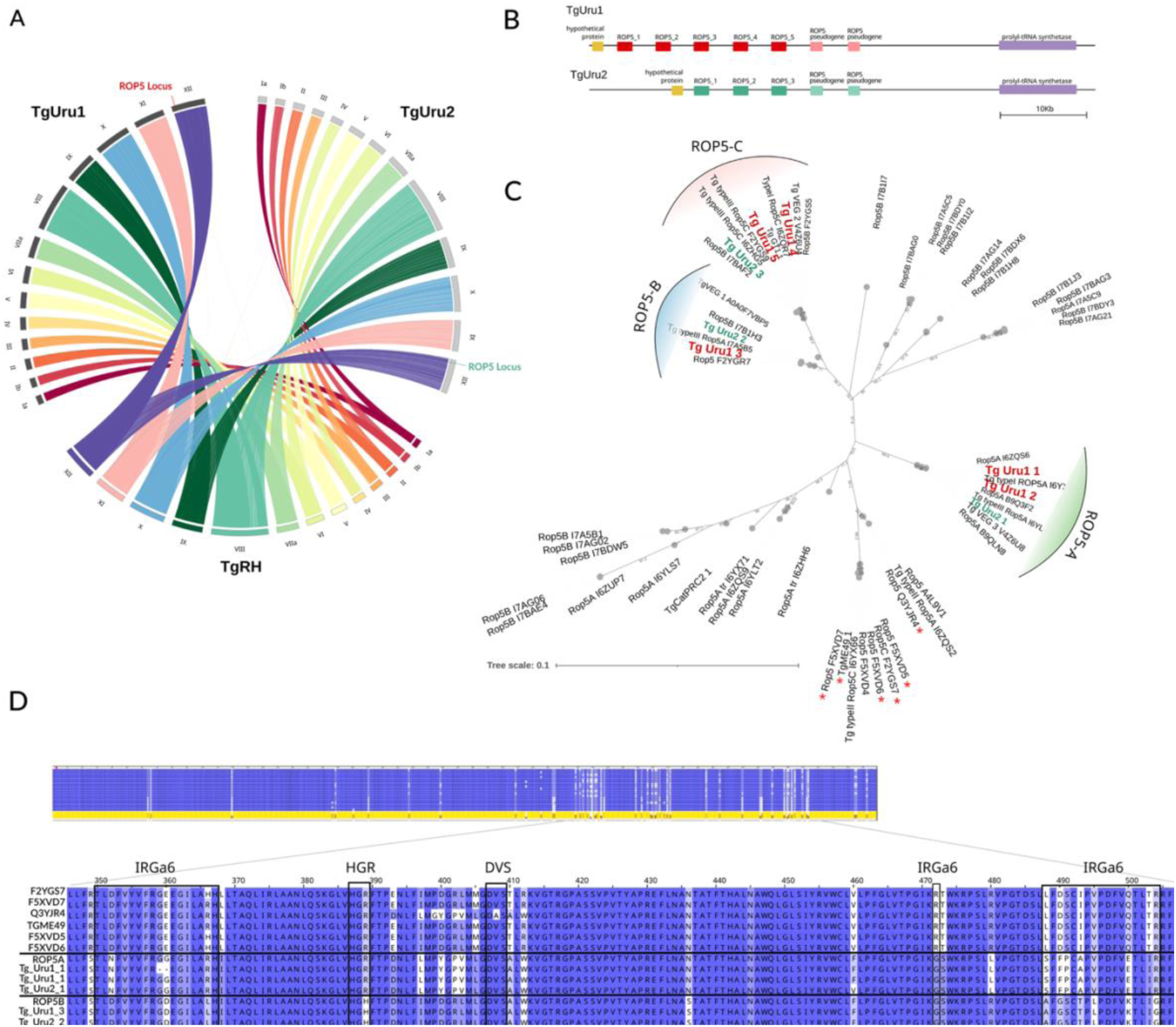
Comparative genomic structure and ROP5 diversity in TgUru1 and TgUru2. **A. Genomic structure.** Circos Plot of the genome alignment against RH reference assembly shows conserved chromosomal organization in both isolates TgUru1 and TgUru2, with no major structural rearrangements detected. Scaffolded assemblies recovered 13 chromosomes with lengths and composition comparable to that of TgRH. The ROP5 locus location is indicated. **B. Scaled representation of the ROP5 locus in TgUru1 and TgUru2**. Copy number variation analysis revealed five ROP5 copies in TgUru1 and three in TgUru2. For reference, the RH strain contains four copies (not shown). **C. Phylogenetic reconstruction of ROP5 copies.** Sequence comparisons among ROP5 copies present in TgUru1 and TgUru2, together with 52 reference ROP5 proteins from UniProt. Both isolates harbor representative member of the three canonical A/B/C subtypes (indicated on the tree). Asterisks denote the sequences selected for the subset alignment shown in panel **D**. **Focused alignment of a representative subset of ROP5 proteins.** Alignment of selected ROP5 subtype members, shown with an asterisked in panel C, together with all copies from TgUru1 and TgUru2. The zoomed region highlights the C-terminal pseudokinase domain, where most sequence variants among ROP5 proteins are concentrated.

To quantify genetic divergence, we identified SNPs and indels in each genome relative to representative strains of the three classical lineages—type I (TgRH), type II (TgME49), and type III (TgVEG). Consistent with its close relationship to type I, TgUru1 showed very few variants relative to TgRH (13,302), whereas TgUru2 displayed a markedly higher number of differences (422,318). In contrast, when compared to the type III strain VEG, TgUru1 accumulated substantially more variants (481,708 vs. 292,817 for TgUru2), reflecting TgUru2’s closer relation to type III. Relative to the type II strain TgME49, TgUru1 and TgUru2 carried 544,472 and 554,941 variants, respectively, reflecting a similarly large genetic distance from type II. Taken together, these results suggest that TgUru2 is genetically closer to Type III, whereas TgUru1 shows greater affinity to type I, consistent with our initial *in silico* PCR-RFLP typing.

We next examined copy number variation in virulence-associated effector gene families using the long-read assemblies. We focused on genes encoding rhoptry and dense-granule proteins involved in host invasion and immune modulation, including the pseudokinases ROP5 and ROP18, as well as additional virulence-associated effectors (GRA24, BFD1, MIC2, GRA7, and GRA17, among others). While the virulent reference strain TgRH carries four copies of ROP5, TgUru1 displays five. Conversely, TgUru2 bears three copies of ROP5 (**Figure 3B**).

ROP5 displays underlying variability, reportedly encompassed by three canonical ROP5 subtypes named A, B, and C. These subtypes have been shown to differ in functionality[55], and virulence has been shown to correlate not only with the number of copies [18], but also with the strain’s ROP5/ROP18 combination[21]. We thus set out to determine the ROP5 subtypes present in each genome. For this, we performed a phylogenetic analysis including all ROP5 gene copies identified in TgUru1 and TgUru2 together with the 52 ROP5 protein sequences available in UniProt (**Figure 3C**). The resulting phylogeny shows that both Uruguayan isolates harbor representatives of the three canonical ROP5 subtypes (A, B, and C). As expected, subtypes B and C grouped more closely to each other than to subtype A.

The tree also revealed substantial sequence diversity within the ROP5 family, extending beyond the three subtypes originally defined by [56] (**Figure 3C**). In several cases, proteins annotated in UniProt as ROP5A, ROP5B, or ROP5C did not cluster with their expected subtype, and in some instances intermingled with other groups, indicating inconsistencies in subtype annotation and highlighting the complexity of the ROP5 paralogous family. This pattern was also evident in the pairwise amino acid identity matrix derived from the multiple sequence alignment (**Supplementary Figure 4**), which clearly distinguished the canonical type A clade from the more closely related Type B/C variants. Overall amino acid identity across all sequences averaged 94.5%, reflecting substantial conservation within each subtype but pronounced divergence between them (**Figure 3D**).

Inspection of the full multiple sequence alignment (**Supplementary Figure 5**) showed that most of the variability is concentrated in the C-terminal pseudokinase domain, between positions 330 and 545. A focused alignment of a representative subset of proteins confirmed this pattern and revealed specific substitutions and small indels in some TgUru1 and TgUru2 copies (**Figure 3D**). Notably, although the subtype-defining polymorphic sites were clearly identifiable, copies assigned to the same subtype (A, B, or C) showed little or no variation within the important regions in the pseudokinase domain (IRGa6, HGR and DVS), except for a two-guanine deletion observed in the TgUru1_1 ROP5 copy.

Beyond ROP5, several effectors involved in immune regulation—including GRA15 and GRA12C—are present in both Uruguayan isolates but absent in RH, suggesting lineage-specific retention or expansion among South American strains. Conversely, genes such as GRA4 and AP2XI-4 are absent in TgUru2, which may reflect localized gene loss, differential pseudogenization, or assembly differences supported by long-read evidence.

Finally, to place the Uruguayan isolates in a broader evolutionary context, we performed a comparative phylogenomic analysis using 100 conserved protein-coding genes across 51 *T. gondii* strains (**Figure 4**). This dataset included 47 hybrid isolates from the Americas described by [51], the previously reported Uruguayan strain CASTELLS, and the canonical laboratory reference strains TgME49, TgVEG, TgRH and TgPRU. Phylogenetic reconstruction revealed that the three Uruguayan strains fall into distinct clades. CASTELLS, previously associated with highly virulent non-clonal lineages, clustered separately and showed resemblance to several non-clonal Brazilian strains, yet remained genetically distinct. TgUru1 grouped with TgRH type I, and other hybrid strains from the Americas and the USA, whereas TgUru2 clustered within a lineage III–associated clade containing strains from the Americas and Africa. TgUru2 showed the closest, though still distant, relationship to TgVEG. None of the Uruguayan isolates clustered with TgME49 or TgPRU. Overall, these results indicate that Uruguayan *T. gondii* strains are highly divergent from one another, reflecting multiple, independent evolutionary origins.

**FIGURE 4.**
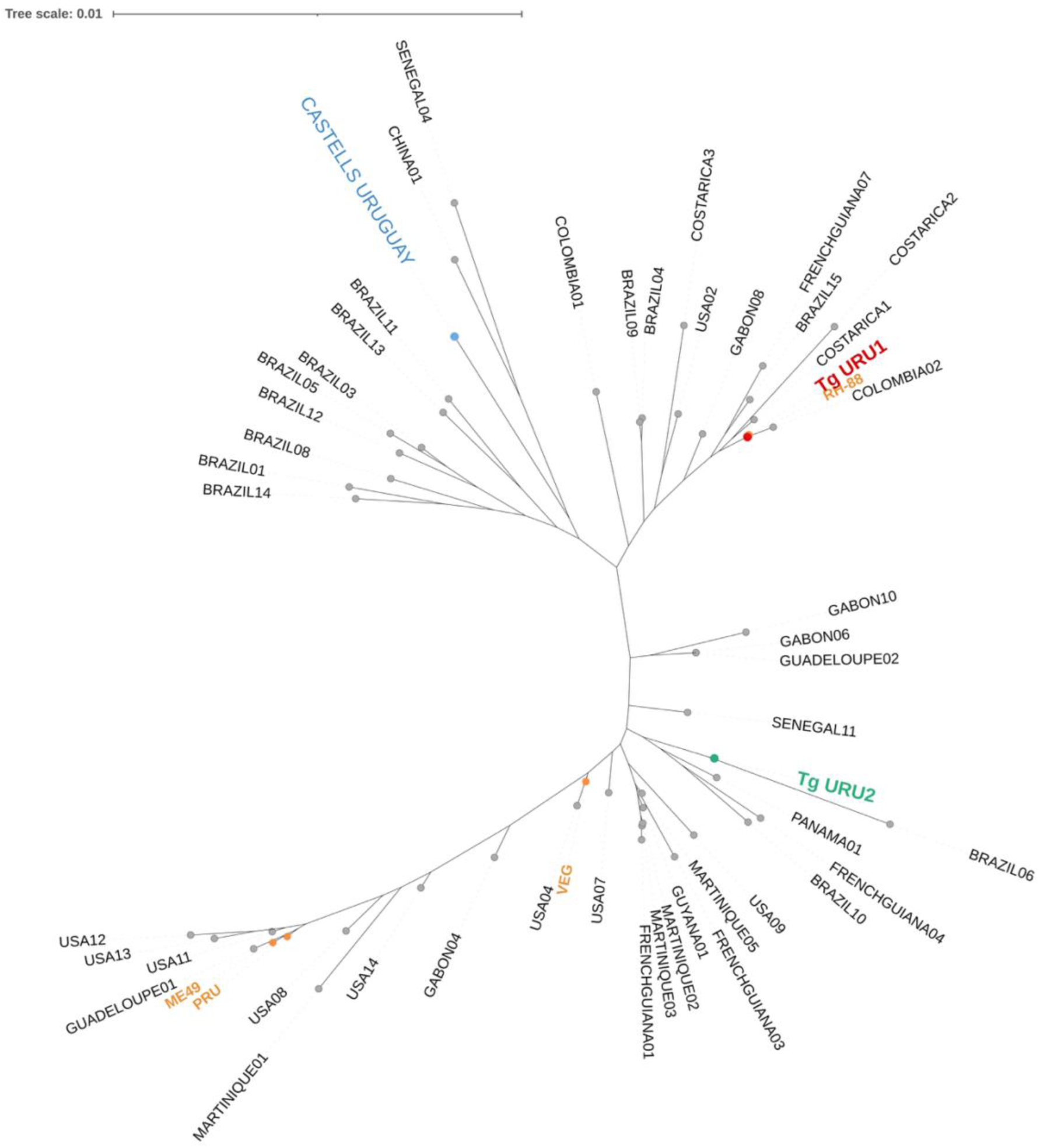
Genome-wide phylogeny of *Toxoplasma gondii* strains using 100 single-copy nuclear genes. Maximum-likelihood phylogeny inferred from 100 single-copy nuclear genes across 51 *Toxoplasma gondii* strains. Three Uruguayan isolates occupy distinct evolutionary positions and do not cluster with TgME49 or TgPRU, reflecting independent origins. CASTELLS groups with non-clonal Brazilian lineages, TgUru1 clusters with type I–related American hybrids, and TgUru2 falls within a lineage III–associated clade.

## DISCUSSION

Toxoplasmosis is considered one of the most globally distributed zoonotic diseases of both human and other animals [57,58]. It is estimated that one-third of the human and animal population has antibodies against *Toxoplasma*, indicating they are either chronically infected or have been exposed to the parasite [58,59]. Sheep are particularly sensitive to infection during pregnancy, and *T. gondii* is a well-recognized leading cause of abortion in this specie worldwide [60]. In Uruguay, records from over 30 years ago identified *T. gondii* as a critically important infectious agent involved in gestational and neonatal losses in sheep [20]. A more recent study analyzing the causes of abortion in 100 ovine cases in Uruguay determined that 27% were still attributable to *T. gondii* infection [22].

Herein, we demonstrated that out of 17 abortion causing strains in sheep, 59% presented non-clonal genotypes with a combination of clonal strain loci and/or presence of previously unidentified RFLP patterns (12%). Though the diversity of *T. gondii* genotypes present in sheep in Uruguay was previously unknown to this resolution; the genetic heterogeneity in the country has been documented in humans. Serotyping studies led to the identification of 36 distinct serotypes, most (73.8%) corresponding to non-clonal variants, with clonal types II and III being rarely detected [61]. Prior work from our group also identified ample variability with molecular resolution in human samples from maternal seroconversion cases [30]. The coexistence of such diverse genetic backgrounds suggests frequent recombination and local diversification events in Uruguay. Conspicuously, the genotypes identified in humans cluster partially with those identified in sheep, suggesting that similar, if not the same, strains are present in both species – consistent with the zoonotic potential of toxoplasmosis. These findings underscore the paradigmatic relevance of *T. gondii* to the *One Health* concept whereby environmental, human and animal infections impact both animal and public health.

An inherent limitation of genotyping *T. gondii* from biological samples, be it by RFLP or by other genomic based analyses, is the dependence on the quality of recovered DNA from samples exposed to adverse environmental conditions and the presence of PCR inhibitors in clinical samples. Herein, we failed to amplify and resolve the full extent of genotypic variability; only being able to unequivocally identify 8 distinct genotypes. This number is likely an underestimation which reflects the inherent limitations of conventional. However, our expanded phylogenetic analysis, which considers SNPs beyond those affecting the restrictions patterns considered by RFLP analysis, revealed no overlapping strains, except for TgShUru15 and 16 which originated from a placenta and its matching fetus respectively. Our data suggests that the 16 remaining samples might correspond to distinct strains. However, all genotypes remained incompletely characterized as there were typing amplicons we were unable to obtain, precluding us from concluding this unambigously. Nonetheless, our findings are consistent with those reported regionally and worldwide. The growing number of epidemiological surveys across diverse regions and host species has revealed extensive genotypic diversity in *T. gondii*, highlighting a complex population structure with strong geographical signatures. Although hundreds of genetic variants—including numerous non-clonal and non-archetypal strains—have been detected, most human infections in Europe and North America still derive largely from the archetypal type II and III lineages [62]. In contrast, South America, considered the probable center of origin for the species [63], harbors an exceptionally diverse set of lineages. Prior work in Brazil, Colombia, and Argentina has shown that human infections in these countries are predominantly caused by non-clonal, non-archetypal strains [13,51,64–67]. Consistent with this regional complexity, the genotypes identified in our study differ from those previously isolated in Argentina by Pardini and colleagues [67], do not match any of the previously reported Brazilian genotypes, and only one genotype overlapped with two previously reported genotypes detected in samples from Central and North America (TgShUru17 with TgCKGa04 and TgSKMs3).

Despite growing recognition of the remarkable genetic diversity of *T. gondii* in South America, the phenotypic consequences of most non-archetypal and recombinant lineages remain largely unknown. Whereas the virulence and pathogenesis of the three clonal types are well-characterized in model systems, the majority of non-clonal and non-archetypal genotypes cannot yet be linked to defined biological features. Herein, we isolated two novel *T. gondii* strains, TgUru1 and TgUru2, which displayed unique non-clonal genotypes consistent with recombinant ancestry. Their phenotypic characterization allowed us to correlate their genotype to relevant physiological properties. Both isolates were recovered from ovine abortion cases, a disease outcome consistent with elevated parasite virulence, yet the isolates exhibited different virulence and growth profiles in mice and cellular culture. TgUru1 showed extreme virulence and rapid intracellular proliferation, whereas TgUru2 displayed moderate virulence but enhanced cystogenic capacity. These contrasting phenotypes among genetically distinct strains illustrate how genetic diversity may influence infection dynamics and potentially the severity of infection when present. Highly virulent genotypes such as TgUru1 could be associated with acute infections leading to early fetal loss, while less virulent but persistent strains such as TgUru2 might underline late term abortions of subclinical gestational infections, or even underlie vertical transmission in cases of infection reactivation – a phenomenon observed to occur in experimentally infected sheep [68]. However, the phenotype observed in the mouse model should be correlated with that in sheep.

Distinct genetic backgrounds have been associated with varying degrees of disease severity in humans. While some strains have been shown to display heightened virulence, reports also describe non-archetypal strains as moderately virulent or non-virulent [13,69–72]. Reports also suggest that parasite diversity may underly placental tropism affecting the parasite’s propensity to be vertically transmitted [73]. In addition, lack of cross-strain protection from vertical transmission during pregnancy has been experimentally shown in sheep [68] and demonstrated to occur naturally in humans [74]. The latter carries implications for development of prophylatic strategies, and challenges our assumptions on transmission, particularly in settings where strain diversity is the norm. Hence, there is a pressing need for systematic parasite isolation and experimental characterization. Expanding the collection of viable South American isolates will be essential to establish robust genotype–phenotype relationships and to fully understand the evolutionary and clinical consequences of this extraordinary population structure.

In light of the phenotypic diversity found for our two isolates, we pursued the identification of differences between the strains, extending beyond their genotyping markers. While overall genomic structure remained unchanged in the isolates, our analysis revealed clear differences in the number of ROP5 copies between TgUru1 and TgUru2. Phylogenetic classification showed that the copies present in both strains fall within the three canonical subtypes (A, B, and C) described by [56]. Despite the broader diversity reported for the ROP5 family in *T. gondii*, the copies assigned to each subtype were virtually identical across their key functional regions—including the IRGa6-binding interface, the HGR/HGH catalytic triad, and the DVS motif—except for a two-guanine deletion detected in one TgUru1 copy. The overall conservation of the pseudokinase domain is consistent with its essential role in modulating the IRG response in mice, a mechanism known to be a major determinant of acute virulence [17,75].

Although previous studies have reported no global correlation between total ROP5 copy number and virulence across the major *T. gondii* lineages [17,56], a link between the expansion of the loci and changes in virulence in admixed strains has been established more recently [18]. The two isolates analysed here share the same ROP5 allele types yet differ both in virulence and in the number of ROP5A and ROP5C copies. Notably, Niedelman and collaborators showed that higher ROP5 expression correlates with reduced IRG coating, supporting a non-enzymatic, dose-dependent role for ROP5 in IRG evasion [76]. Within this framework, variation in ROP5 copy number could modulate the effective dosage of IRG-inhibitory ROP5 variants when allelic background is held constant. In this context, the higher number of ROP5 copies observed in TgUru1 may partially contribute to its increased virulence relative to TgUru2.

Although sequence identity among ROP5 proteins was high (∼95%), our phylogeny confirms that the ROP5 repertoire extends beyond the three original subtypes defined from laboratory strains [56]. The incorporation of more allelic sequences from diverse geographical origins has revealed a continuum of ROP5 diversity, much of it concentrated in the C-terminal pseudokinase region where this family evolves under strong positive diversifying selection [17,56]. This expanded variability likely reflects adaptation to different host IRG repertoires, an idea supported by the patterns observed in South American strains and by previous analyses showing that positively selected residues tend to localize on the IRGa6-binding surface of ROP5 [17].

Our detailed phylogenomic analysis shows that the three Uruguayan isolates (TgUru1, TgUru2 and the previously obtained CASTELLS) fall into distinct clades, consistent with independent evolutionary histories. This pattern mirrors the genomic structure described by Galal et al. (2022), where chromosome-painting ancestry profiles correspond closely to genome-wide phylogenetic relationships [51]. Alongside with previous work showing that South American *T. gondii* populations are highly admixed and often display mosaic genomes shaped by limited recombination, these results underscore the remarkable genetic diversity characteristic of “New World” parasites [51,77]. Notably, despite the much larger number of Brazilian genomes included in our analysis, the breadth of diversity represented by only three Uruguayan isolates is comparable, suggesting that Uruguay harbors a similarly heterogeneous reservoir of *T. gondii* genotypes. The absence of major geographic barriers between Uruguay and Brazil—combined with the documented movement of wildlife and livestock (and people) across the dry border—likely contributes to this shared diversity, a pattern also observed in other zoonotic pathogens such as *Mycobacterium bovis* [78]. A more comprehensive assessment of how geographical barriers influence the regional distribution of *T. gondii* genotypes will require broader genomic sampling, especially of underrepresented regions like Argentina, Paraguay, and Chile. Overall, our findings reveal a high genetic and phenotypic diversity among *T. gondii* strains associated with ovine abortion in Uruguay, highlighting the circulation of non-clonal lineages with probable links to human infections. Our findings underscore the importance of regional genomic surveillance to better understand transmission dynamics, pathogenic potential, and zoonotic risk in South America.

## ACKNOWLEDGEMENTS

LB, PF, FG, MEF are researchers of PEDECIBA and the national system of researchers (SNI-ANII). We are deeply thankful for the everlasting contributions of Dr. Alvaro Freyre, and Dr. Dario Falcon, to the *Toxoplasma* research in Uruguay upon which we have built on in this manuscript.

## DATA AVAILABILITY

The datasets generated during and/or analyzed during the current study are available in the NCBI’s GenBank database, and can be retrieved using the individual identifiers provided in supplementary material

## AUTHOR CONTRIBUTIONS

L.T.-H. contributed to methodology, investigation, data curation, formal analysis, and writing – original draft. P.F., A.M.C., and A.V. contributed to methodology and investigation, with P.F. also contributing to formal analysis. M.D. and S.F. provided resources. F.G. contributed to conceptualization and resources. L.B. and M.E.F. contributed to conceptualization, investigation, data curation, formal analysis, writing – review & editing, and supervision. M.E.F. additionally contributed to project administration and funding acquisition. All authors read and approved the final manuscript

## ADDITIONAL INFORMATION

The authors declare no competing financial and non-financial interest

